# Synchronized EEG with two galvanically-separated miniature wireless behind-the-ear EEG sensors

**DOI:** 10.1101/2025.03.15.643275

**Authors:** Ruochen Ding, Simon Geirnaert, Charles Hovine, Alexander Bertrand

**Affiliations:** KU Leuven, Department of Electrical Engineering (ESAT), STADIUS Center for Dynamical Systems, Signal Processing and Data Analytics and with Leuven.AI – KU Leuven institute for AI; KU Leuven, Department of Neurosciences, Research Group ExpORL

## Abstract

We present a wireless EEG sensor network consisting of two miniature, wireless, behind-the-ear sensor nodes with a size of 2 cm *×* 3 cm, each containing a 4-channel EEG amplifier and a wireless radio. Each sensor operates independently, each having its own sampling clock, wireless radio, and local reference electrode, with full electrical isolation from the other. The absence of a wire between the two nodes enhances discreetness and flexibility in deployment, improves miniaturization potential, and reduces wire artifacts. A third identical node acts as a USB dongle, which receives and synchronizes the data from the two behind-the-ear nodes. The latter allows to process the 2 *×* 4 channel EEG as if all 8 channels are sampled synchronously, allowing the use of signal processing algorithms that exploit inter-channel correlations. To demonstrate this synchronized processing, we recorded auditory steady-state responses (ASSRs) at both ears and processed them with data-driven multi-channel filters to optimize the ASSR signal-to-noise ratio, demonstrating a more reliable ASSR detection compared to a single-ear setup.

## 1. INTRODUCTION

Electroencephalography (EEG) is a widely used non-invasive technique for recording brain activity, with applications spanning clinical diagnostics, neuroscience research, and brain-computer interfaces (BCIs) [1]. It has been extensively applied in areas such as epilepsy monitoring, sleep studies, mental health assessment, and auditory attention decoding [2], [3], [4], [5]. Traditional EEG systems rely on bulky equipment, extensive wiring, and large electrode arrays. These limitations restrict their portability and usability in everyday settings, which leads to a growing interest in wearable EEG systems designed for comfortable, long-term brain monitoring in real-life environments.

Mobile head-mounted EEG systems have been developed to improve the portability of traditional EEG setups by integrating portable amplifiers and wireless data transmission [6], [7]. While these systems mitigate some issues of traditional setups, their bulky design make them uncomfortable for long-term daily use. Furthermore, the physical wiring across the headset introduces electrode-displacement artifacts or wire artifacts.

This research is funded by the Research Foundation Flanders (FWO) project No G081722N, junior postdoctoral fellowship fundamental research of the FWO (for S. Geirnaert, No. 1242524N), the European Research Council (ERC) under the European Union’s research and innovation program (grant agreement No 802895 and No 101138304), Internal Funds KU Leuven (project IDN/23/006), and the Flemish Government (AI Research Program). Views and opinions expressed are however those of the author(s) only and do not necessarily reflect those of the European Union or the granting authorities. Neither the European Union nor the granting authorities can be held responsible for them.

Ear electroencephalography (ear-EEG) has emerged as an effective wearable EEG method that takes advantage of the anatomical structure around the ear [8]. This design provides stable electrode placement and reduces user discomfort, which allows for continuous brain monitoring in everyday environments and serves as a more practical alternative to traditional scalp EEG systems.

Recent advancements in ear-EEG technology have led to the development of in-ear and behind-the-ear electrode grids. In-ear systems place electrodes within the ear canal, providing high concealment and improved comfort for long-term use [9]. These systems have been validated for applications such as sleep stage monitoring and real-time detection of brain activity anomalies, including seizure patterns [3], [8], [10]. A detailed summary of in-ear EEG systems is provided in [11]. However, their small form factor often limits the number of available channels, which reduces the spatial resolution compared to head-mounted systems [12]. Behindthe-ear solutions, such as the cEEGrid, employ flexible electrode arrays positioned around the ear to improve comfort and reduce visibility [13]. The development, applications, and limitations of cEEGrid technology are reviewed in [14]. These ear-EEG systems have shown effectiveness in recording continuous EEG, event-related potentials, and neural oscillations. An overview of various ear-EEG technologies is provided in [15].

However, existing ear-EEG systems still face several limitations. Most of them rely on long wires (between both ears and/or towards a centralized amplifier), which not only reduce the system’s discreetness but also introduce challenges such as motion artifacts caused by cable movement or electrode displacement. Additionally, the presence of long cables increases susceptibility to environmental interference, including electromagnetic noise from power lines and surrounding electronic devices, further degrading signal quality.

To address these challenges, we propose a miniaturized 2ear system consisting of two compact wireless sensor nodes with a size of 2 cm *×* 3 cm, each containing a 4-channel EEG amplifier and a wireless radio. A distinctive feature, absent in previous ear-based systems, is that both nodes operate independently without physical wiring between the two ears, each using a different reference electrode. This miniaturized design and absence of long wires reduces susceptibility to power line noise, and wire-pulling/motion artifacts. The independent operation of each sensor enhances system reliability by allowing one sensor to continue functioning even if the other fails. It also simplifies maintenance and replacement, improving overall practicality. A tailored synchronization protocol between the (independently) digitized sensor signals maintains temporal alignment, enabling accurate multichannel EEG acquisition, allowing the use of signal processing algorithms that exploit the spatial correlation structure between all 8 channels.

We validate the system through auditory steady-state response (ASSR) experiments, demonstrating its ability to capture synchronized, high-quality EEG signals. ASSRs are periodic brain responses evoked by periodic auditory stimuli such as a sinusoidally amplitude-modulated carrier signal. These ASSRs are phase-locked to the modulation-frequency of the stimulus [16]. The results confirmed reliable detection of ASSRs at both ears simultaneously. Notably, using optimal data-driven filtering which combines the channels of both ears allows to achieve better and more robust ASSR detection compared to single-ear setups.

## II METHODS

### A. Sensor node design and functionality

Our system consists of two miniature sensor nodes placed behind the left and right ear and one data sink node connected to a computer via USB. Each sensor node consists of a 2 cm *×* 3 cm double-sided six-layer PCB, to which 4 (+1 reference) electrodes can be connected, allowing to record 4 EEG channels per sensor (we refer to [17] for more details on the PCB and firmware design). The two behindthe-ear sensor nodes continuously transmit their EEG data to the data sink node, which synchronizes the data streams and forwards the aligned data to the computer for further analysis. The system is powered by 250 mAh Li-Po batteries, allowing up to 5 hours of real-time EEG data streaming to the laptop. Fig. 1 shows an initial prototype, illustrating how the sensor nodes, battery, and electrodes could be positioned behind both ears of a subject, e.g., using a 3D-printed C-shape holder (to enhance concealment in future iterations, the electrode wires could be integrated into the C-shape holder, similar to the cEEGrid [14]).

**Fig. 1:**
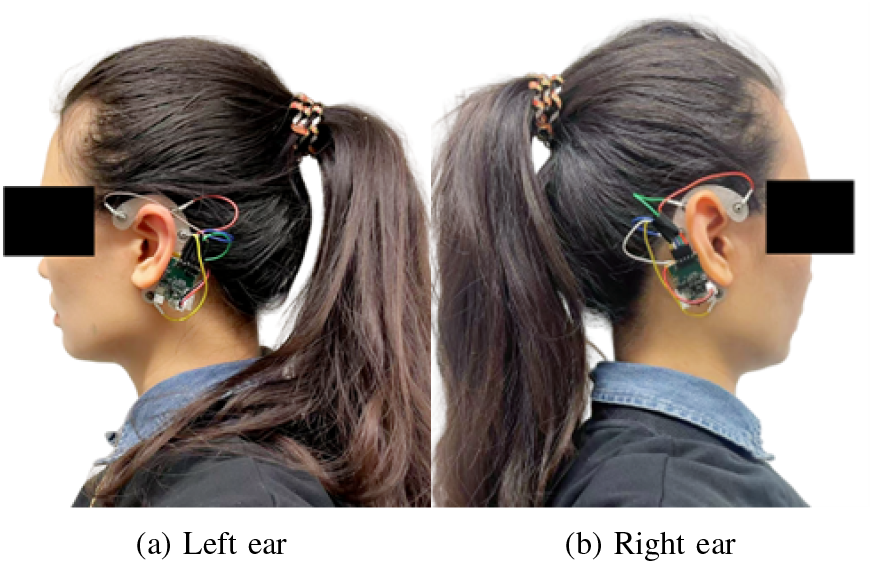
Prototype displaying the two behind-the-ear wireless sensor nodes. Note: this is a mock-up used for illustration. In the experiments described in Section III, the electrodes were directly affixed to the skin using adhesive tape, without the use of the 3D-printed C-shaped holder (*These photographs depict the author Ruochen Ding*).

The PCB integrates three core modules: (1) an Analog Front-End (AFE) for high-resolution EEG signal acquisition, (2) an integrated Microcontroller Unit (MCU) with a built-in radio module for data processing and wireless transmission, and (3) a power management module for efficient power distribution (Fig. 2). Bias resistors replace the traditional right leg drive (RLD) circuit, reducing the number of required electrodes while maintaining signal quality without the need for a dedicated bias or ground electrode. A summary of the key specifications of the sensor node is provided in Table I.

**Fig. 2:**
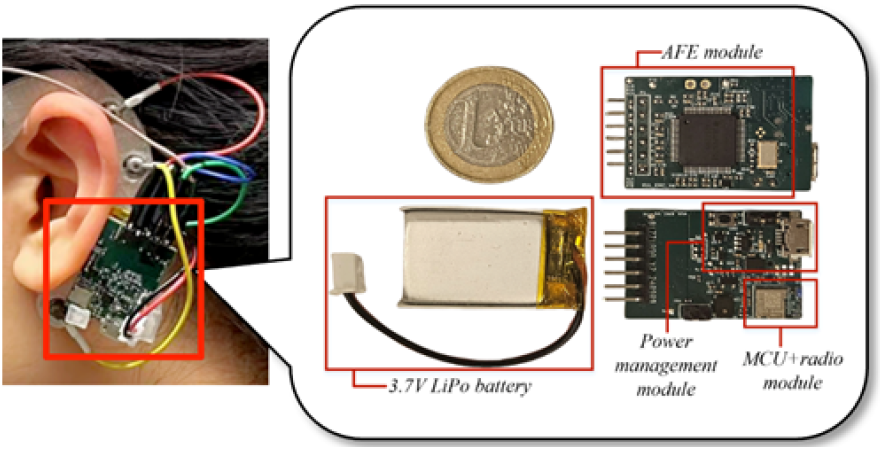
Front and back view of the PCB.*(This photograph depicts the author, Ruochen Ding)*

**TABLE 1:**
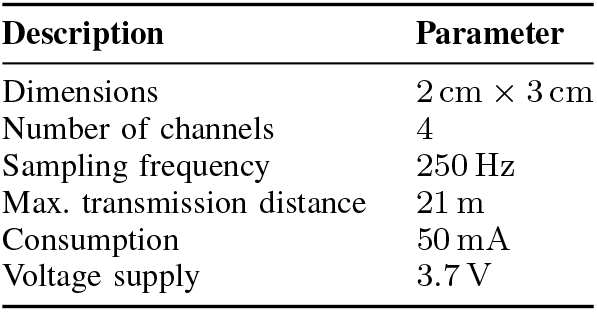
Parameters of the EEG sensor node.

The lack of a connecting wire between the two nodes prevents the use of a shared reference electrode. As a result, each node has its own local reference electrode. Combined with the miniaturized design, this setup restricts measurements to local EEG potentials using short inter-electrode distances.

However, we will demonstrate that despite this limitation, the EEG signals from both ears can still exhibit common components, which can be leveraged in multi-channel signal processing pipelines.

Another consequence of the absence of a wire between the nodes is that each node must use its own local sampling clock to digitize the data. Time synchronization between the EEG data streams from both nodes is achieved through a protocol similar to the Precision Time Protocol (PTP), which dynamically compensates for clock drift by estimating wireless link latency and adjusting timestamps. This functionality has been validated in prior work, which confirmed its effectiveness in maintaining long-term sample-level synchronization [17].

### B. Spatial filtering for enhanced signal quality

Our goal is to combine the EEG data from both sensors in a synchronous multi-channel signal processing setup for detecting ASSRs—periodic brain responses evoked by an auditory stimulus that is amplitude-modulated at a specific modulation frequency *f*_*m*_. To this end, we will employ optimal data-driven max-SNR filtering as in [16], [18]. We start from the following data model:

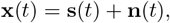

with **x**(*t*) ∈ ℝ^*C×*1^ the recorded *C*-channel EEG signals consisting of the periodic ASSR **s**(*t*) at the modulation frequency *f*_*m*_ and uncorrelated background EEG noise **n**(*t*). In our experimental setup *C* can be either 8 (both ears) or 4 (single ear).

To find the spatial filter 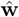 that optimally combines the different EEG channels to enhance the ASSR-SNR in a datadriven way, we optimize the SNR:

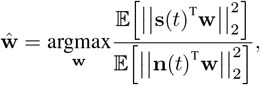

where 𝔼 denotes the expectation operator. Using spatial covariance matrices **R**_*s*_ = 𝔼 **s**(*t*)**s**(*t*)^T^ *∈* ℝ^*C×C*^ (ASSR) and **R**_*n*_ = 𝔼 [**n**(*t*)**n**(*t*)^T^] *∈* ℝ^*C×C*^ (noise), this boils down to:

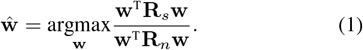

However, the ASSR **s**(*t*) and, therefore, signal covariance matrix **R**_*s*_, are not known, making it impossible to solve this optimization problem. Nonetheless, we can find the same optimal spatial filter 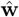 by optimizing the (signal+noise)-to-noise ratio instead of the signal-to-noise ratio in (1):

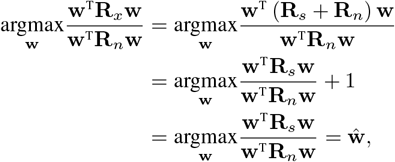

where the first equality holds because the ASSR and noise are assumed to be uncorrelated, such that **R**_*x*_ = **R**_*s*_ + **R**_*n*_. The covariance matrices **R**_*x*_ (EEG, containing both ASSR and noise) and **R**_*n*_ (noise) can be estimated based on the same recording by taking different frequency ranges assuming coherent noise in a small band around the modulation frequency, i.e., [*f*_*m*_ − *δ, f*_*m*_ +*δ*] for **R**_*x*_ (containing the ASSR and noise) and [*f*_*m*_ Δ, *f*_*m*_ *δ*] [*f*_*m*_ + *δ, f*_*m*_ + Δ] for **R**_*n*_ (containing only noise), with bandwidths *δ ≪*Δ [16].

Equation (1) can now be solved by taking the generalized eigenvector corresponding to the largest generalized eigenvalue of the generalized eigenvalue decomposition (GEVD) of matrix pencil (**R**_*x*_, **R**_*n*_) [19]. Using the estimated spatial filter 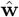, the output signal 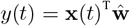 can be computed, on which ASSR detection can be performed (see Section III-B).

## III EXPERIMENTAL SETUP AND RESULTS

### A. Measurement setup

To provoke ASSRs, participants were presented with binaural broadband CE-chirp stimuli which were amplitude-modulated at a rate of 40 Hz [20] and played through headphones at a sampling rate of 48 kHz at a comfortable volume. EEG recordings were made using the proposed two-node behind-the-ear system, using standard Ag/AgCl electrodes that were affixed behind the ear with skin-friendly tape. Before placement, the skin was prepared with Nuprep gel to remove dead skin layers and oil. Parker Signa Gel was applied to the electrodes to reduce impedance and ensure optimal electrical contact with the skin. The experiment involved six participants, each completing three trials of 3 minutes each, resulting in a total of 9 minutes of EEG data per subject. One subject participated twice (Subject 3 = Subject 6) in two different sessions on different days, such that there are 7 recordings (here noted as 7 subjects) in total. Our experiment was approved by the KU Leuven Social and Societal Ethics Committee.

### B. ASSR detection

For ASSR detection, we employed the data-driven max-SNR filtering approach explained in Section II-B. Every 3-minute trial was split into three 1-minute segments, over which cross-validation was performed. Per trial, the GEVD-based spatial filter was thus trained on two 1-min segments and tested on the left-out one. This was repeated for all three segments in each 3-min trial, and all 3-min trials. The bandpass filters extracting the EEG samples to populate the covariance matrices (**R**_*x*_, **R**_*n*_) are estimated using secondorder sinc filters (i.e., twice applying an ideal sinc filter) with *δ* = 0.125 Hz and Δ = 6 Hz. The analysis was conducted using three configurations: using all four channels from the left node only, the right node only, or the combined 8-channel data from both nodes.

The SNR based on the single-channel output signal *y*(*t*) was computed per 1 s-window for each 1-min segment (generated with 50% overlap) to evaluate the system’s ability to detect ASSRs at 40 Hz. This SNR is defined as the energy at the modulation frequency *f*_*m*_ over the energy in a small band of *±*6 Hz around the modulation frequency, excluding the modulation frequency itself. Furthermore, an F-test with *α*-level = 0.01 is conducted per 1 s-window to check whether a significant ASSR is detected or not [21].

Fig. 3 presents the resulting average SNRs across 1 swindows per 1-min segment for every subject under the different sensor configurations, as well as an average across all subjects. Combining EEG signals from both sensor nodes typically results in an SNR level that is at least as high as the best single-ear filter, and often even better, both on a per-subject basis and on average across subjects. This means that using the two sensors at both ears results in a more robust ASSR detection than when only using one node at one ear. This is confirmed by the detection accuracy (i.e., the number of significant ASSR detections across 1 s-windows), which is 89.9% when using both sensors, versus 70.4% (left ear)/89.3% (right ear) when using only one sensor.

**Fig. 3:**
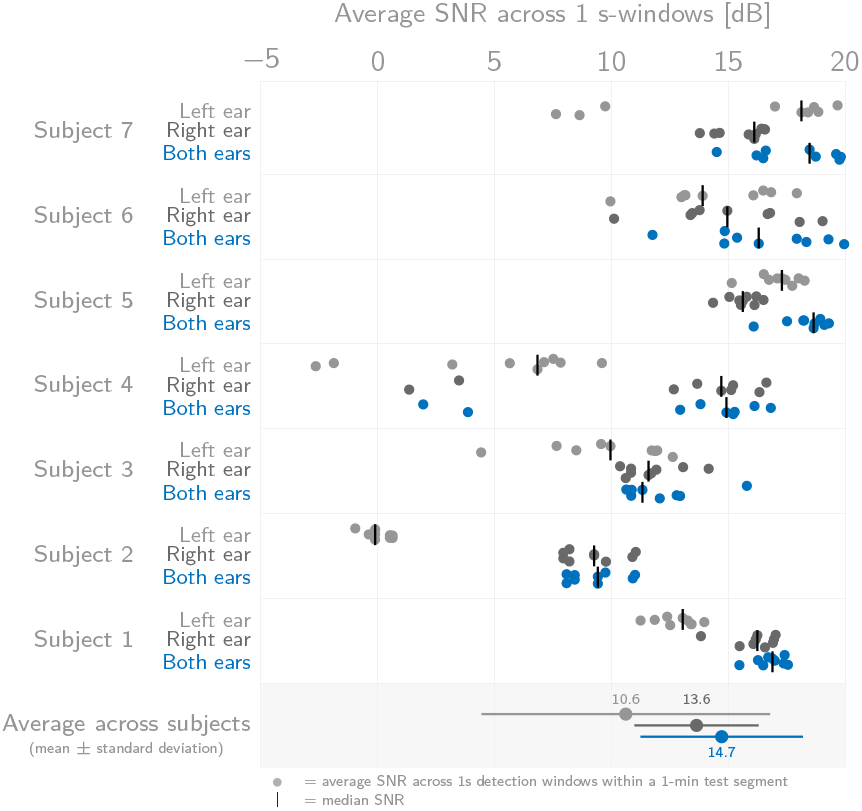
The average SNR across 1 s-windows is consistently as good or better using both sensors at both ears w.r.t. to using only one sensor at one ear.

This improved and more robust ASSR detection using two wirelessly connected EEG sensor nodes would have only be possible with a successful synchronization between the data streams of both sensors. Only then the (mere) spatial filtering of all 8 channels could result in a beneficial effect w.r.t. the SNR and detection accuracy, as otherwise out-of-phase summation would harm ASSR detection. Therefore, the successful ASSR detection results showcase the ability of using the proposed wirelessly synchronized miniature two-node sensor system for recording ear-EEG in auditory neuroscience applications.

## IV. CONCLUSION

We have introduced a compact, wireless, and synchronized 2-node behind-the-ear EEG system that effectively captures synchronized, multi-channel EEG signals. ASSR experiments confirmed the system’s ability to reliably detect auditory responses, which demonstrated its suitability for ear-EEG applications in auditory neuroscience and continuous brain monitoring. The wireless, modular design facilitates a flexible deployment, where the absence of wires between the nodes reduces the susceptibility to motion or wire(-pulling) artifacts. Despite the fact that both nodes sample their data independently, a synchronization protocol allows that their EEG channels can be jointly processed in a data-driven multi-channel signal processing pipeline, as demonstrated here via SNR-optimal ASSR detection. Future work will focus on further reducing the system’s size and power consumption and incorporating real-time processing to expand its potential for advanced ear-EEG applications and continuous brain monitoring in daily life.

## ACKNOWLEDGMENTS

The authors thank Anna Sergeeva for providing the broadband chirp stimuli.

## Footnotes

1 Ruochen Ding, Simon Geirnaert, Charles Hovine, and Alexander Bertrand are with KU Leuven, Department of Electrical Engineering (ESAT), STADIUS Center for Dynamical Systems, Signal Processing and Data Analytics and with Leuven.AI – KU Leuven institute for AI.

2 Simon Geirnaert is also with KU Leuven, Department of Neurosciences, Research Group ExpORL.

